# Periodic attention operates faster during more complex visual search

**DOI:** 10.1101/2021.09.22.460906

**Authors:** Garance Merholz, Laetitia Grabot, Rufin VanRullen, Laura Dugué

## Abstract

Attention has been found to sample visual information periodically, in a wide range of frequencies below 20 Hz. This periodicity may be supported by brain oscillations at corresponding frequencies. We propose that part of the discrepancy in periodic frequencies observed in the literature is due to differences in attentional demands, resulting from heterogeneity in tasks performed. To test this hypothesis, we used visual search and manipulated task complexity, i.e., target discriminability (high, medium, low) and number of distractors (set size), while electro-encephalography was simultaneously recorded. We replicated previous results showing that the phase of pre-stimulus low-frequency oscillations predicts search performance. Crucially, such effects were observed at increasing frequencies within the theta-alpha range (6-18 Hz) for decreasing target discriminability. In medium and low discriminability conditions, correct responses were further associated with higher post-stimulus phase-locking than incorrect ones, in increasing frequency and latency. Finally, the larger the set size, the later the post-stimulus effect peaked. Together, these results suggest that increased complexity (lower discriminability or larger set size) requires more attentional cycles to perform the task, partially explaining discrepancies between reports of attentional sampling. Low-frequency oscillations structure the temporal dynamics of neural activity and aid top-down, attentional control for efficient visual processing.

## Introduction

Covert attention selects task-relevant stimuli among the constant flow of sensory information (in the absence of eye and head movement) to facilitate their processing [1-3]. A large literature (for review [4-8]) shows that spatial attention samples visual information periodically at low frequencies, i.e., < 20 Hz [9-19] and that this sampling is supported by brain oscillations at corresponding frequencies [20-24]. Critically, the frequencies reported in the literature show fairly large discrepancies (Table 1), which to date remains understudied. One parameter that may be involved is attentional demand, as it was shown to correlate with neural activity in posterior and frontal areas [44], visual neuronal activity [45], and with individual alpha peak frequency [46]. Here, we explicitly tested whether the variability in frequencies reported across the attentional sampling literature may be due to varying attentional demands, potentially caused by the disparity of tasks.

**Table 1.**
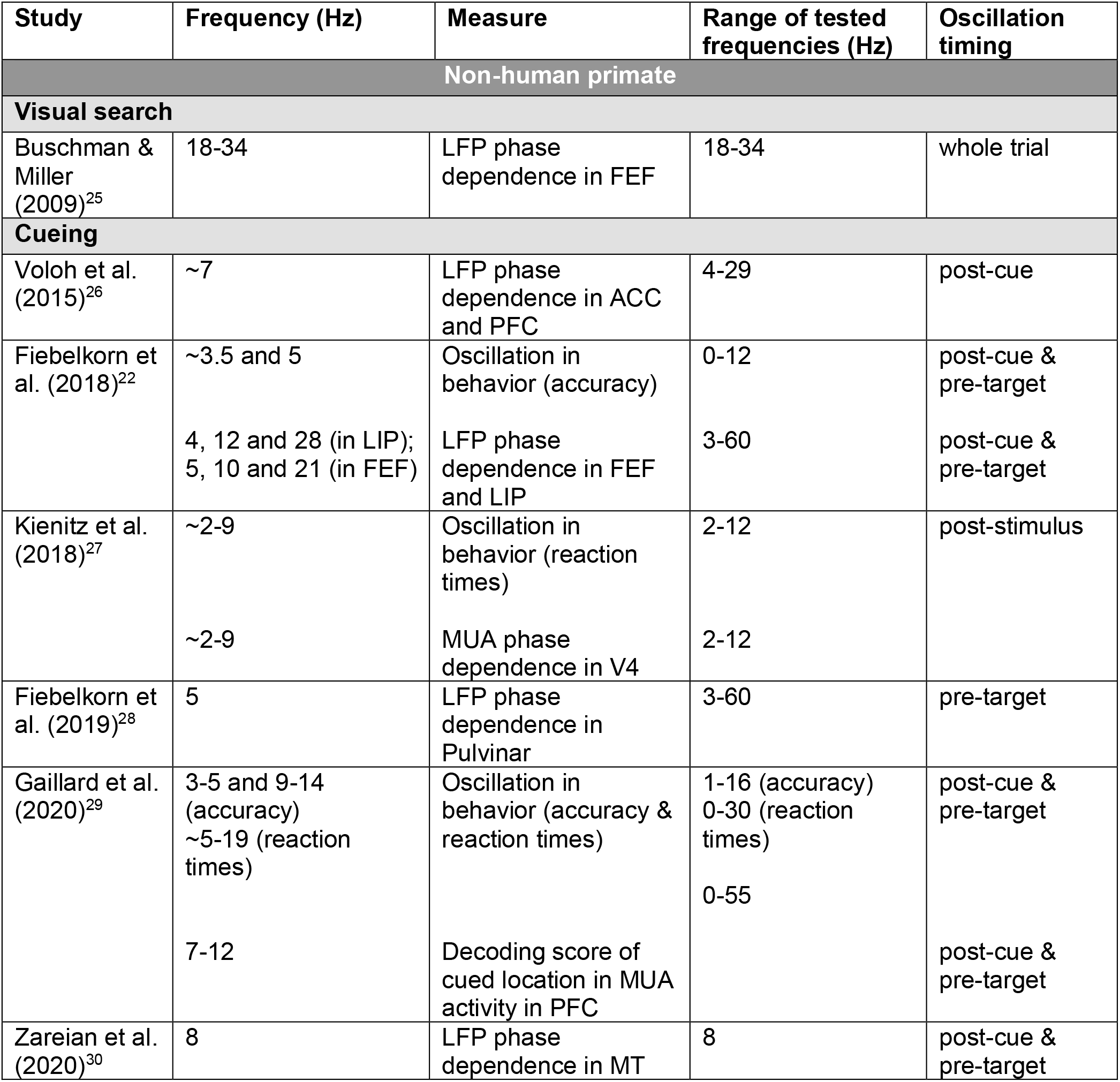
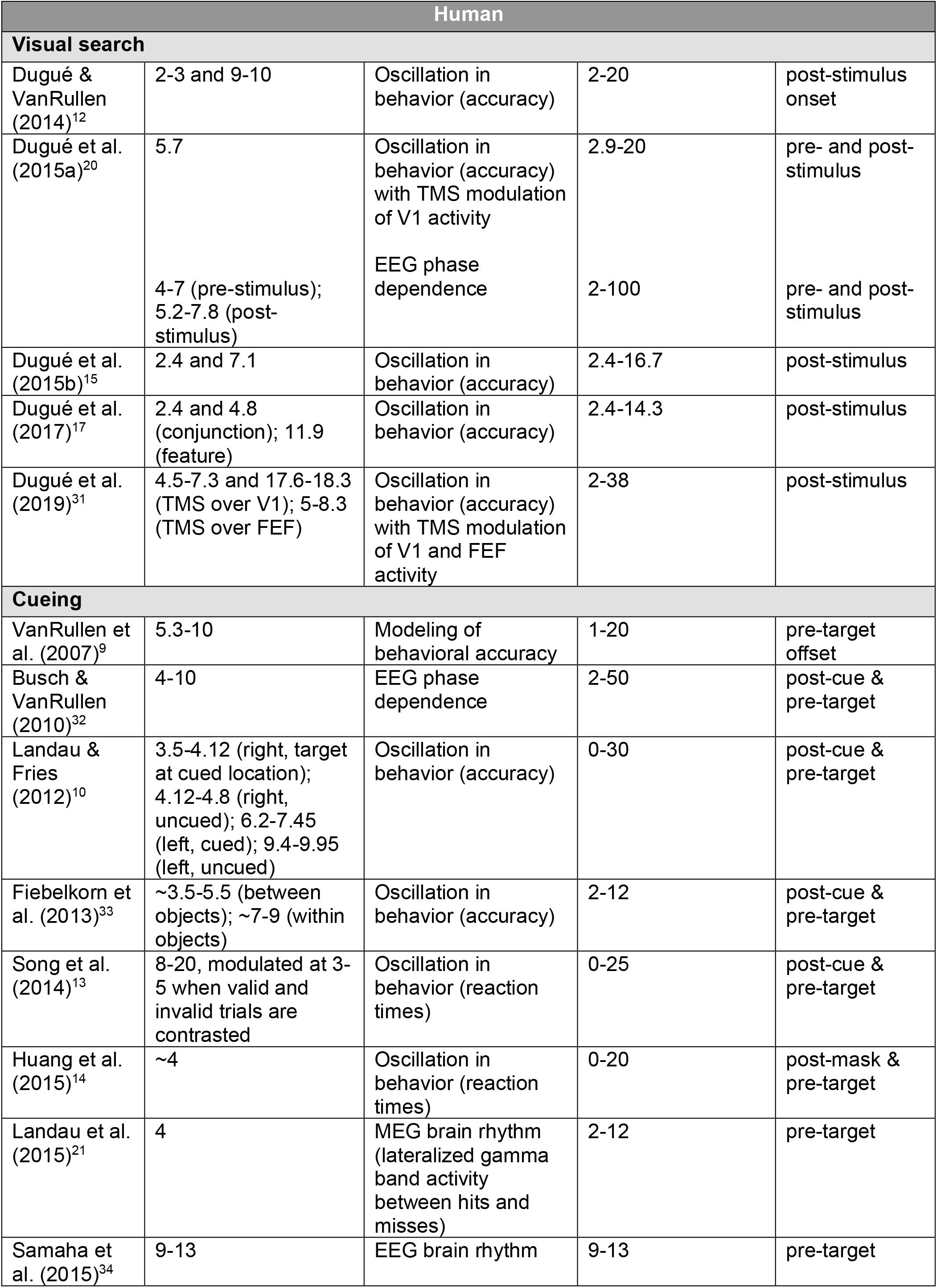
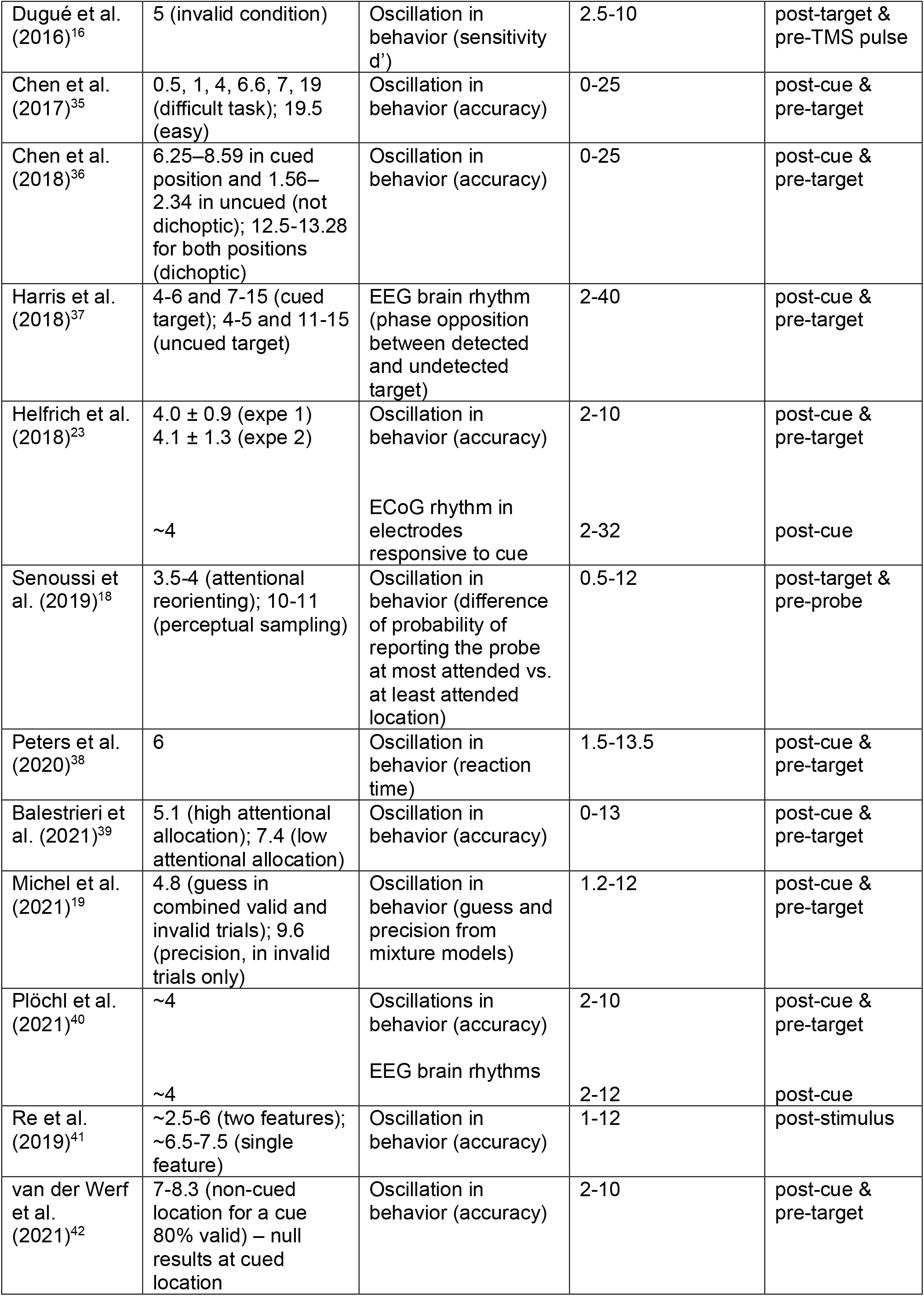
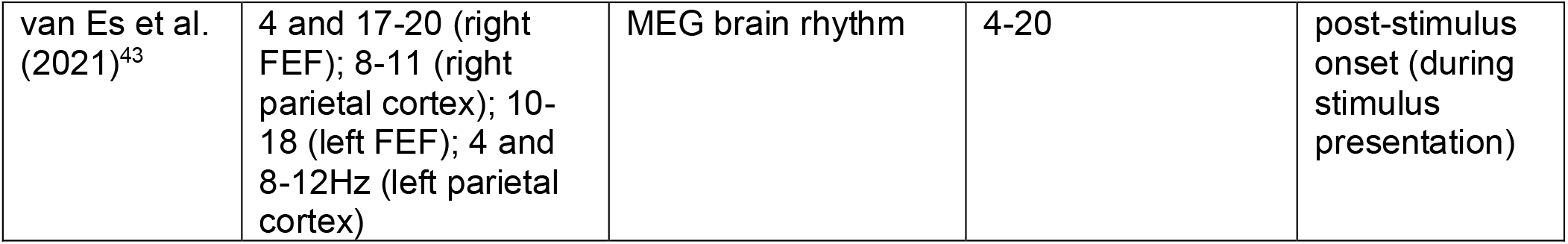
Studies demonstrating periodic phase effects during covert visual attention tasks in human and non-human primates. These studies were the “Attention and search” studies in VanRullen (2018)^96^ and updated based on a literature search (until January 2022). The same criteria were used to select the studies, i.e., studies explicitly manipulating covert visual attention and showing a link between instantaneous phase and behavioral performance (from these studies, we only report results regarding oscillations in behavior, as well as analyses linking instantaneous neural phase and behavioral performance). Studies “entraining” brain oscillations were not included (see VanRullen, 2018^96^, for more details). Properties of the periodicity are organized by species (non-human primate or human) and nature of the attentional manipulation (visual search or cueing; no other manipulation was found). Within these fields, the following sub-fields are included, from left to right: Study (chronological order); Frequency of significant rhythm (some of these values were not clearly reported in the publication and thus correspond to estimations based on Figure reading; indicated with ∼); Measure used in the study to reveal periodicity; Range of all tested frequencies; Oscillation timing relative to trial events. Note that the table does not include negative findings. LPF: Local Field Potential. FEF: Frontal Eye Field. ACC: Anterior Cingulate Cortex. PFC: Lateral Prefrontal Cortex. LIP: Lateral Intraparietal Area. MUA: Multi-Unit Activity.

| Study | Frequency (Hz) | Measure | Range of tested frequencies (Hz) | Oscillation timing |
| --- | --- | --- | --- | --- |
| **Non-human primate** |  |  |  |  |
| **Visual search** |  |  |  |  |
| Buschman & Miller (2009)<sup>25</sup> | 18-34 | LFP phase dependence in FEF | 18-34 | whole trial |
| **Cueing** |  |  |  |  |
| Voloh et al. (2015)<sup>26</sup> | ~7 | LFP phase dependence in ACC and PFC | 4-29 | post-cue |
| Fiebelkorn et al. (2018)<sup>22</sup> | ~3.5 and 5 | Oscillation in behavior (accuracy) | 0-12 | post-cue & pre-target |
|  | 4, 12 and 28 (in LIP);<br>5, 10 and 21 (in FEF) | LFP phase dependence in FEF and LIP | 3-60 | post-cue & pre-target |
| Kienitz et al. (2018)<sup>27</sup> | ~2-9 | Oscillation in behavior (reaction times) | 2-12 | post-stimulus |
|  | ~2-9 | MUA phase dependence in V4 | 2-12 |  |
| Fiebelkorn et al. (2019)<sup>28</sup> | 5 | LFP phase dependence in Pulvinar | 3-60 | pre-target |
| Gaillard et al. (2020)<sup>29</sup> | 3-5 and 9-14 (accuracy)<br>~5-19 (reaction times) | Oscillation in behavior (accuracy & reaction times) | 1-16 (accuracy)<br>0-30 (reaction times)<br>0-55 | post-cue & pre-target |
|  | 7-12 | Decoding score of cued location in MUA activity in PFC |  | post-cue & pre-target |
| Zareian et al. (2020)<sup>30</sup> | 8 | LFP phase dependence in MT | 8 | post-cue & pre-target |

**Table 1. Studies demonstrating periodic phase effects during covert visual attention tasks in human and non-human primates.**
| Human |  |  |  |  |
| --- | --- | --- | --- | --- |
| Visual search |  |  |  |  |
| Dugué & VanRullen (2014)<sup>12</sup> | 2-3 and 9-10 | Oscillation in behavior (accuracy) | 2-20 | post-stimulus onset |
| Dugué et al. (2015a)<sup>20</sup> | 5.7<br>4-7 (pre-stimulus);<br>5.2-7.8 (post-stimulus) | Oscillation in behavior (accuracy) with TMS modulation of V1 activity<br>EEG phase dependence | 2.9-20<br>2-100 | pre- and post-stimulus<br>pre- and post-stimulus |
| Dugué et al. (2015b)<sup>15</sup> | 2.4 and 7.1 | Oscillation in behavior (accuracy) | 2.4-16.7 | post-stimulus |
| Dugué et al. (2017)<sup>17</sup> | 2.4 and 4.8 (conjunction); 11.9 (feature) | Oscillation in behavior (accuracy) | 2.4-14.3 | post-stimulus |
| Dugué et al. (2019)<sup>31</sup> | 4.5-7.3 and 17.6-18.3 (TMS over V1); 5-8.3 (TMS over FEF) | Oscillation in behavior (accuracy) with TMS modulation of V1 and FEF activity | 2-38 | post-stimulus |
| Cueing |  |  |  |  |
| VanRullen et al. (2007)<sup>9</sup> | 5.3-10 | Modeling of behavioral accuracy | 1-20 | pre-target offset |
| Busch & VanRullen (2010)<sup>32</sup> | 4-10 | EEG phase dependence | 2-50 | post-cue & pre-target |
| Landau & Fries (2012)<sup>10</sup> | 3.5-4.12 (right, target at cued location); 4.12-4.8 (right, uncued); 6.2-7.45 (left, cued); 9.4-9.95 (left, uncued) | Oscillation in behavior (accuracy) | 0-30 | post-cue & pre-target |
| Fiebelkorn et al. (2013)<sup>33</sup> | ~3.5-5.5 (between objects); ~7-9 (within objects) | Oscillation in behavior (accuracy) | 2-12 | post-cue & pre-target |
| Song et al. (2014)<sup>13</sup> | 8-20, modulated at 3-5 when valid and invalid trials are contrasted | Oscillation in behavior (reaction times) | 0-25 | post-cue & pre-target |
| Huang et al. (2015)<sup>14</sup> | ~4 | Oscillation in behavior (reaction times) | 0-20 | post-mask & pre-target |
| Landau et al. (2015)<sup>21</sup> | 4 | MEG brain rhythm (lateralized gamma band activity between hits and misses) | 2-12 | pre-target |
| Samaha et al. (2015)<sup>34</sup> | 9-13 | EEG brain rhythm | 9-13 | pre-target |
| Dugué et al. (2016)<sup>16</sup> | 5 (invalid condition) | Oscillation in behavior (sensitivity d') | 2.5-10 | post-target & pre-TMS pulse |
| Chen et al. (2017)<sup>35</sup> | 0.5, 1, 4, 6.6, 7, 19 (difficult task); 19.5 (easy) | Oscillation in behavior (accuracy) | 0-25 | post-cue & pre-target |
| Chen et al. (2018)<sup>36</sup> | 6.25–8.59 in cued position and 1.56–2.34 in uncued (not dichoptic); 12.5-13.28 for both positions (dichoptic) | Oscillation in behavior (accuracy) | 0-25 | post-cue & pre-target |
| Harris et al. (2018)<sup>37</sup> | 4-6 and 7-15 (cued target); 4-5 and 11-15 (uncued target) | EEG brain rhythm (phase opposition between detected and undetected target) | 2-40 | post-cue & pre-target |
| Helfrich et al. (2018)<sup>23</sup> | 4.0 ± 0.9 (expe 1)<br>4.1 ± 1.3 (expe 2) | Oscillation in behavior (accuracy) | 2-10 | post-cue & pre-target |
|  | ~4 | ECoG rhythm in electrodes responsive to cue | 2-32 | post-cue |
| Senoussi et al. (2019)<sup>18</sup> | 3.5-4 (attentional reorienting); 10-11 (perceptual sampling) | Oscillation in behavior (difference of probability of reporting the probe at most attended vs. at least attended location) | 0.5-12 | post-target & pre-probe |
| Peters et al. (2020)<sup>38</sup> | 6 | Oscillation in behavior (reaction time) | 1.5-13.5 | post-cue & pre-target |
| Balestrieri et al. (2021)<sup>39</sup> | 5.1 (high attentional allocation); 7.4 (low attentional allocation) | Oscillation in behavior (accuracy) | 0-13 | post-cue & pre-target |
| Michel et al. (2021)<sup>19</sup> | 4.8 (guess in combined valid and invalid trials); 9.6 (precision, in invalid trials only) | Oscillation in behavior (guess and precision from mixture models) | 1.2-12 | post-cue & pre-target |
| Plöchl et al. (2021)<sup>40</sup> | ~4 | Oscillations in behavior (accuracy) | 2-10 | post-cue & pre-target |
|  | ~4 | EEG brain rhythms | 2-12 | post-cue |
| Re et al. (2019)<sup>41</sup> | ~2.5-6 (two features); ~6.5-7.5 (single feature) | Oscillation in behavior (accuracy) | 1-12 | post-stimulus |
| van der Werf et al. (2021)<sup>42</sup> | 7-8.3 (non-cued location for a cue 80% valid) – null results at cued location | Oscillation in behavior (accuracy) | 2-10 | post-cue & pre-target |
| van Es et al. (2021)<sup>43</sup> | 4 and 17-20 (right FEF); 8-11 (right parietal cortex); 10-18 (left FEF); 4 and 8-12Hz (left parietal cortex) | MEG brain rhythm | 4-20 | post-stimulus onset (during stimulus presentation) |

Visual search tasks – looking for a target among distractors – have been largely used to study attention (for review [47,48]), and more specifically its temporal dynamics. A periodic attentional mechanism implies cycles of discrete sampling windows over time. Visual search is particularly adapted to test the link between these attentional cycles, neural oscillations and task demands, because task demands may be easily manipulated by varying target discriminability among distractors or varying the number of distractors.

It has been proposed that attentional sampling during visual search tasks relies on periodic neural activity [5,7,20,49,31]. Particularly, it was shown that the pre- and post-stimulus phase of theta oscillations (4-8 Hz) distinguished correct from incorrect trials [20]. Interestingly, when disrupting low-level occipital (V1) or high-level attentional (FEF) areas in humans using transcranial magnetic stimulation (TMS), performance was modulated periodically at low frequencies (< 20 Hz) [20,49,31]. In other words, specific delays at which TMS was administered during the task led to increased behavioral performance while others led to decreased performance, with preferred and unpreferred delays alternating periodically at low frequencies. This result was interpreted as evidence that the sampling process of attentional selection during visual search is periodic and is supported by brain oscillations at low frequencies. It is also consistent with hierarchical models of visual search, which posit that effortful search involves iterative communication between occipital and frontal areas [50-53]. Such an iterative process may be subtended by corresponding low-frequency neural oscillations [5,7,20,49,31,54].

Here, we recorded electro-encephalography (EEG) during visual search tasks with varying attentional load to test the link between brain oscillations and task demands, and further characterize periodic attentional sampling. We first replicated previous results showing that pre-stimulus phase [20,32,55-63] and post-stimulus phase-locking across trials [20,64,65] of low-frequency oscillations predict successful task performance (same task as in [20]). Specifically, the phase of spontaneous oscillations (before stimuli onset) predicts trial outcome; and post-stimulus phase alignment (phase-locking) aids performance, which we call phase reset here to refer, strictly, to the mathematical description of the effect [66].

The novelty in the present study lies in the implementation of a gradient of task demands during visual search in two separate experiments. Crucially, we tested the prediction that an increase in the number of cycles required to perform the search would result in an increase in the frequency of the underlying neural oscillations, and/or in a delayed peak of the neural response. Specifically, we manipulated the discriminability of the target (Experiment 1) and the number of items in the search array, i.e., set size (Experiment 2). Our working hypothesis was that during one attentional cycle only a limited amount of information from the search array is processed - either because not all stimuli are attended at once, or because attention is divided among all stimuli and thus less resources are attributed to each stimulus (here, we do not aim to disentangle these two options). It follows that given a fixed search duration, the more complex the search task, the higher the frequency of the neural oscillation required for successful performance - allowing more cycles to be completed in the same amount of time. Moreover, alpha/theta phase realignment has been associated with more efficient sensory processing of targets [20,30]. Therefore, if lower discriminability and additional items entail more attentional cycles, and if phase-locking aids performance while processing the target, a delayed phase reset can be expected, to allow for a periodic sampling process to cycle through the search array and complete the processing of the target.

In Experiment 1, the target and distractors’ shapes were manipulated to obtain three levels of discriminability (high, medium and low, with equalized task performance; blocked). We tested two predictions: (1) the lower the discriminability, the higher the frequency of the pre- and post-stimulus brain oscillations underlying attentional performance; (2) the lower the discriminability and the later the optimal phase reset, resulting in delayed post-stimulus temporal dynamics. Indeed, some phase resetting is to be expected simply due to stimulus onset [66,67], but a stronger reset should lead to better performance [20], hence higher phase-locking is expected in correct trials. Furthermore, if the search requires on average more attentional cycles, as hypothesized when increasing search complexity, then this would translate in an optimal phase reset that appears at a delayed latency.

In Experiment 2, the set size (number of items in the search array, 4 or 8; interleaved) was manipulated. Here, we tested a last prediction: (3) the optimal phase reset affecting behavior occurs at a delayed latency and higher frequency when more items are presented (i.e., higher task complexity). Because the participant cannot predict the set size of the upcoming trial, we did not expect an effect of set size on pre-stimulus activity and thus did not test it – there is no reason to assume here that participants can adapt their sampling frequency in advance.

Altogether, the results confirm our three predictions and suggest that increased attentional demands requires more attentional cycles to perform the task (higher frequency), partially explaining discrepancies between reports of attentional sampling.

## Results

### Behavior

Participants performed a visual search task with varying parameters while electroencephalography (EEG) was simultaneously recorded (Fig. 1). In Experiment 1, the effect of discriminability between target and distractors was tested with three conditions (high, medium, and low discriminability; blocked). We titrated the stimulus-onset asynchrony (SOA) for each participant to ensure that their percentage of correct responses remained around 70% and to minimize task difficulty differences between conditions (mean SOA ± SD: high = 40 ± 5 ms, medium= 82 ± 13 ms, low = 271 ± 26 ms; Fig. 1B). Participants’ performance across all conditions was on average at 66% (SD = 6%, ranging from 55 to 85%, Fig. 1C). For the high discriminability condition, average performance was at 68 ± 8%, 68 ± 9% for medium, and 61 ± 6% for low discriminability. A one-way repeated-measures ANOVA revealed that the percentage of correct responses in Experiment 1 significantly differed between discriminability conditions (F(1,2) = 3.73, p = 0.034, η^2^ = 0.18). Post-hoc 2-by-2 t-tests revealed that performance in the low discriminability condition was significantly lower than in the high and medium discriminability conditions (see Statistics Table 1). There was no significant difference in performance between the high and medium conditions.

**Figure 1.**
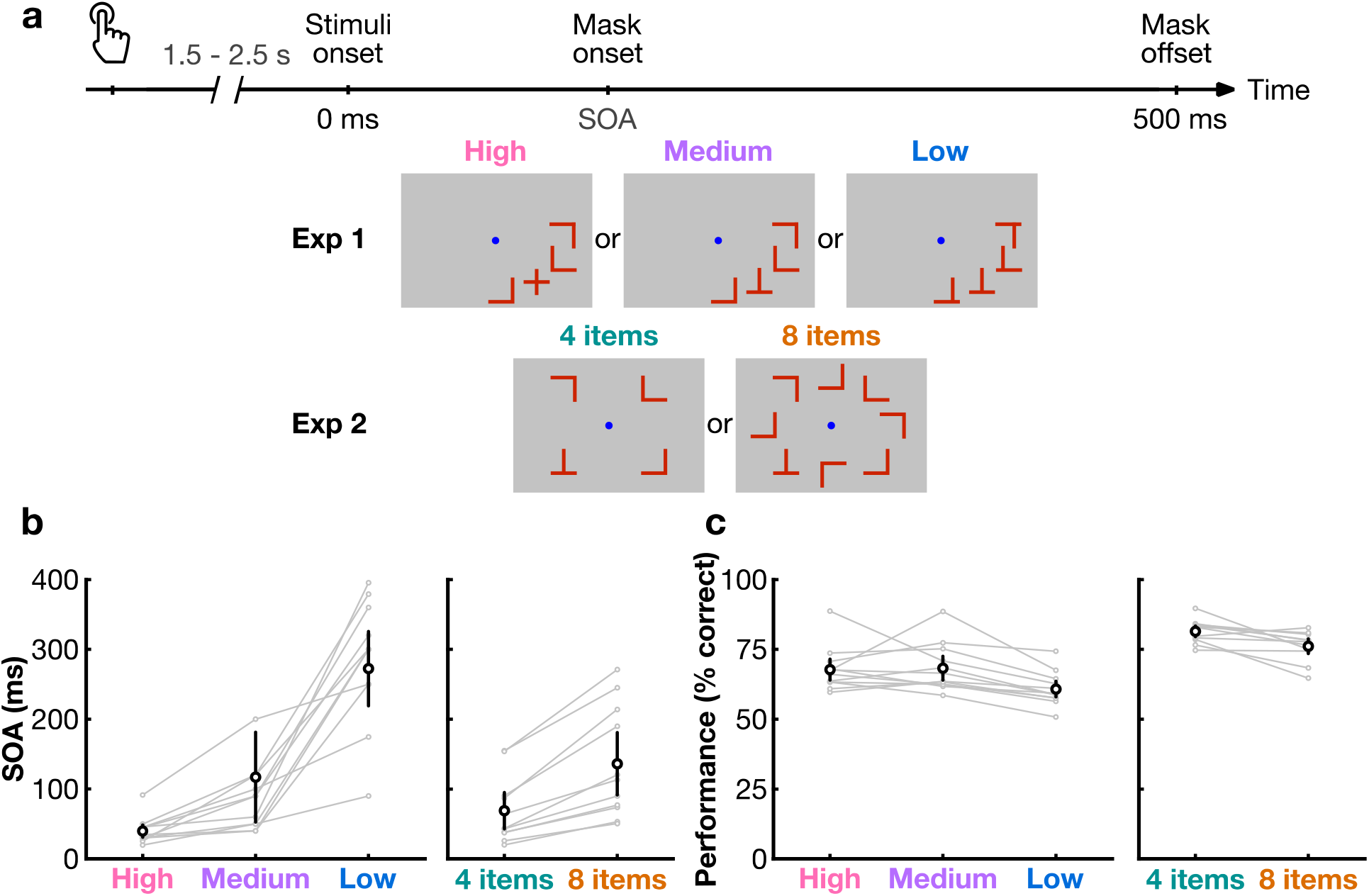
Task design and behavioral results. (**a**) A button press initiated by the participant began the trial. After a random duration between 1.5 and 2.5 seconds, the search array appea red. In Experiment 1, the item shape was manipulated (blocked) to yield high (the target is a +), medium and low (the target is a T) discriminability conditions (always 4 items). In Experiment 2, the number of items was manipulated (4 or 8, interleaved; the target is always a T). The items remained onscreen for the duration of the participant’s titrated SOA, immediately followed by masks, which disappeared after 500 ms from stimuli onset. After mask offset, the participant pressed a button to indicate whether the target was present or absent, ending the trial. The central fixation dot was always present. (**b**) Stimuli duration (SOA) for Experiment 1 (left) and Experiment 2 (right) for each condition. (**c**) Performance expressed as the percentage of correct responses (detecting the presence or absence of the target). Black circles represent the group average performance. Error bars represent the 95% confidence interval. Gray traces show results for individual participants.

**Statistics Table 1.** Performance comparisons (2-by-2 t-tests) between experimental conditions. *: significant p values. CI: 95% Confidence Interval. df: degrees of freedom.

| Condition comparison | *p* | CI | *t* | df | Cohen's *d* |
| --- | --- | --- | --- | --- | --- |
| high vs. low discriminability | < 0.01* | [2.1, 11.9] | 3.11 | 11 | 0.98 |
| medium vs. low discriminability | < 0.001* | [4.3, 10.7] | 5.13 | 11 | 0.98 |
| high vs. medium discriminability | 0.854 | [-6.3, 5.3] | -0.19 | 11 | -0.06 |
| 4 vs. 8 items | 0.008* | [1.7, 9.0] | 3.27 | 10 | 1.06 |

In Experiment 2, the effect of set size was tested with either four or eight items presented (interleaved) in the search array (medium discriminability items were used). The SOA was titrated to minimize task difficulty differences between conditions (mean SOA ± SD: 4 items = 69 ± 43 ms, 8 items = 129 ± 78 ms, Fig. 1B). The average performance across both conditions was at 79% (SD = 4, ranging from 65 to 90 %, Fig. 1C). For the 4-items condition, average performance was at 81 ± 4%, and 76 ± 5% for 8 items. A two-tailed t-test showed that performance was significantly lower in the 8-items condition compared to the 4-items condition (see Statistics Table 1). The medium discriminability condition in Experiment 1 and the 4-items condition in Experiment 2 exhibited the same items and set size, and a two-tailed heteroscedastic t-test revealed that their SOAs were comparable (p = 0.189). However, their performance levels were significantly different (p < 0.0001), perhaps due to inter-individual differences or to item placement on the screen (Fig. 1A).

### Pre-stimulus phase impacts performance at increasing frequency with lower discriminability

We first tested the prediction that the lower the discriminability, the higher the frequency of the pre-stimulus brain oscillations underlying attentional performance. Phase opposition sum (POS) [20,62,68,69] values were estimated for each frequency, pre-stimulus time point and discriminability condition in Experiment 1. A positive POS indicates that the phase is locked in correct trials and that it is locked in the opposite direction in incorrect trials (see Materials and Methods). An FDR-corrected permutation test showed a significant phase opposition between correct and incorrect trials at low frequencies (i.e., 6.6-14.3 Hz; Fig. 2) for the three discriminability conditions. We observed that the peak frequency of POS rose with difficulty: 7 Hz for high, 9.7 Hz for medium and 10.6 Hz for low discriminability condition. The frequency band containing significant POS (blue outline in Fig. 2) had no overlap between the high discriminability condition (6.6 to 7.1 Hz) and the lower-discriminability conditions (medium: 8.5 to 11.1 Hz, low: 8.5 to 14.3 Hz). The topographies at POS peaks (marked by stars) showed that the effect was localized in the occipital and frontal poles (Fig. 2, bottom).

**Figure 2.**
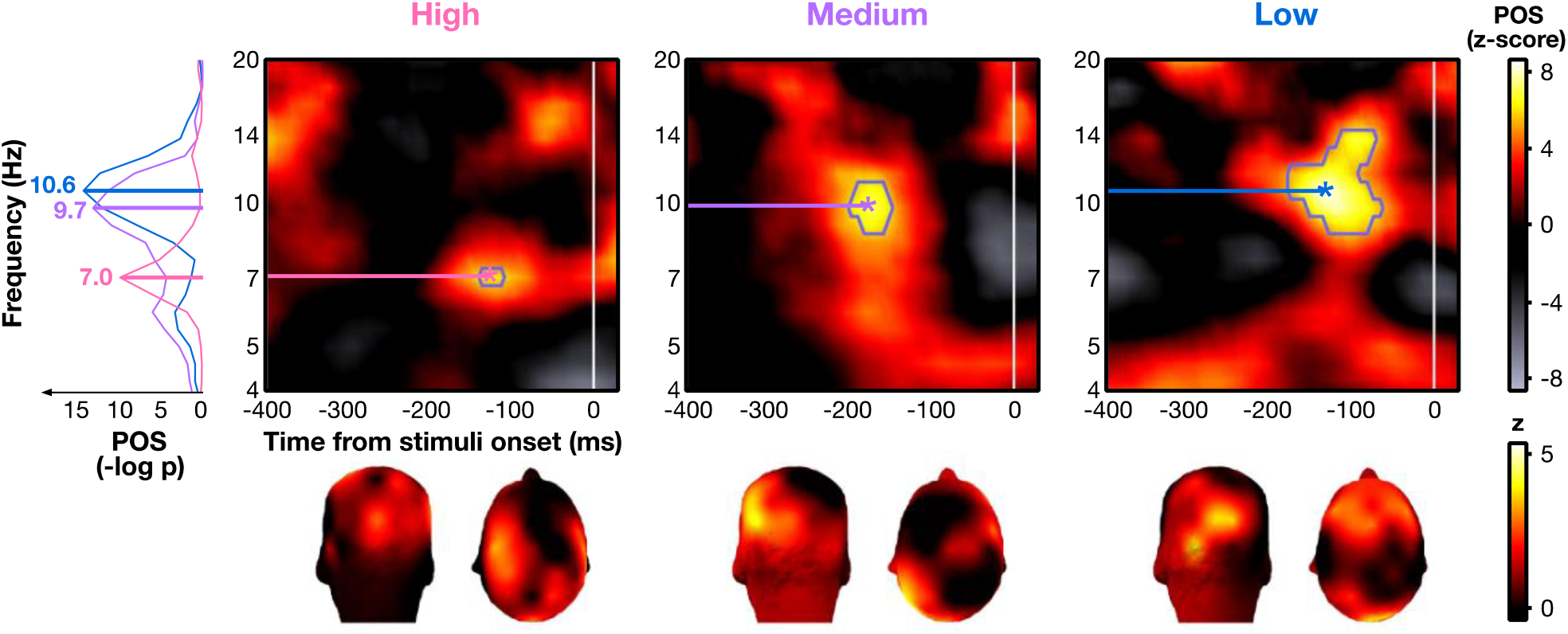
Pre-stimulus phase opposition between correct and incorrect trials in Experiment 1. Z-score of pre-stimulus phase opposition sum (POS) of correct and incorrect trials for the high, medium and low discriminability conditions, combined across all electrodes and all participants. Blue contours indicate areas above the FDR threshold (alpha = 10^−7^, corresponding to p value thresholds of 2.2e-10, 5.0e-10, and 4.7e-9 respectively). Stars indicate the time-frequency point of maximum significance (high: z_max_ = 6.42 at f_max_ = 6.96 Hz and t_max_ = -129 ms; medium: z_max_ = 7.49 at f_max_ = 9.75 Hz and t_max_ = -176 ms; low: z_max_ = 8.91 at f_max_ = 10.60 Hz and t_max_ = -137 ms). Topomaps represent the average POS (z-score) at each electrode across participants for the point of maximum significance, indicated by a star. The left panel plot shows the converted POS (-log p values) of the high, medium and low discriminability conditions (pink, purple and blue traces, respectively) at the timepoint with the maximum z-score (bootstrap-corrected POS in -log(p) of 10.17 for high, 13.48 for medium and 14.68 for low).

To ensure that the observed phase opposition effects were not spuriously generated by amplitude differences, we performed a similar analysis on amplitude. Although we observed significant differences between correct and incorrect trials (high: 8 to 12 Hz and -70 to 0 ms, medium: 6 to 17 Hz and -400 to -130 ms, low: 14 to 18 Hz at -200 ms, see Supplementary Material), these clusters did not overlap with those found for POS (Fig. 2) in the high and low discriminability conditions; the amplitude effects are thus unlikely to be merely due to amplitude differences. Note that there is partial overlap in the medium condition; we thus cannot completely rule out this interpretation for this condition.

### Post-stimulus phase-locking impacts performance only in complex searches

We tested the prediction that the lower the discriminability, the later the optimal reset of neural oscillations involved in attentional search, resulting in slower post-stimulus temporal dynamics. Post-stimulus phase-locking was compared between correct and incorrect trials. The analysis was restricted to the task-relevant frequency band found in the pre-stimulus analysis (from the lowest to the highest FDR-corrected significant frequencies, i.e., 6.6 to 14.3 Hz; Fig. 2). At each time point within each participant and condition, phase-locking difference (PLD) between correct and incorrect trials was averaged across the frequency band of interest. FDR-corrected permutation tests showed that the PLD was significantly different from zero only in lower-discriminability searches (medium: p < 0.001, 82 to 106 ms; low: p < 0.01, 184 to 270 ms), suggesting that correct trials were more phase-locked than incorrect trials in these conditions (Fig. 3A). PLD was not significantly different from zero in the high discriminability condition (p-values remained above the FDR threshold constrained by an alpha parameter of 0.01). The topographies represent the distribution of the activity at the point of maximum significance over all electrodes (Fig. 3A). Note that the effect in the medium condition is localized in fronto-occipital areas, while it is rather parieto-occipital in the low discriminability condition.

**Figure 3.**
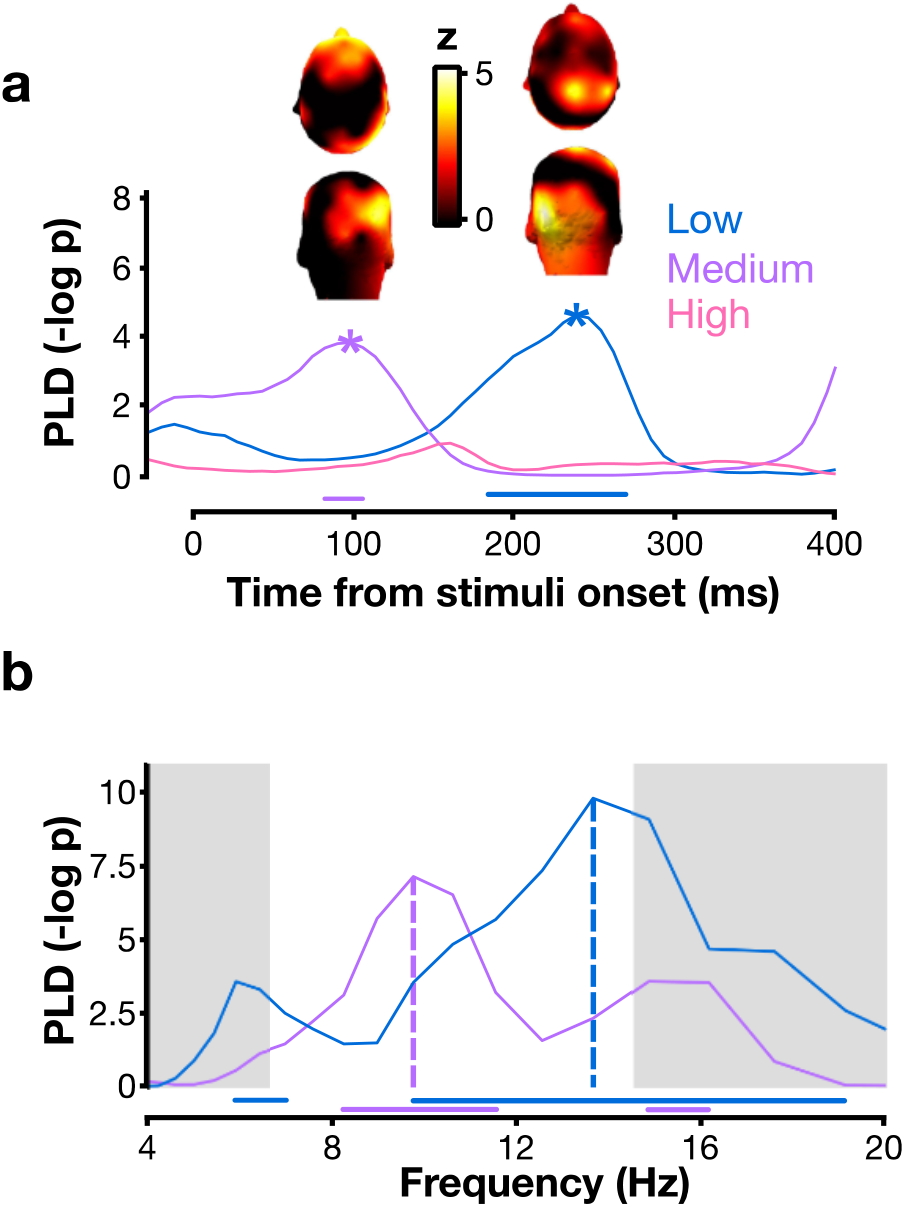
Post-stimulus phase-locking difference between correct and incorrect trials in Experiment 1. (**a**) The phase-locking difference (PLD) is the difference in inter-trial phase coherence (ITC) between correct and incorrect trials. The PLD curves are plotted after averaging across the frequency range of the significant clusters found in pre-stimulus phase opposition (Fig. 2). Stars mark the timepoint of maximum significance on each trace (medium: 98 ms; low: 238 ms). Topomaps represent the PLD at each electrode averaged across participants at the timepoint of maximum significance. (**b**) PLD curves plotted at the timepoints of maximum significance against frequency for the medium (purple) and low (blue) discriminability conditions. The white area indicates the frequency range of interest, taken from the pre-stimulus phase opposition effect. On both panels, horizontal bars along the bottom indicate that the PLD of the corresponding curve (blue: low discriminability; purple: medium discriminability) is above the significance threshold corrected for multiple comparisons (FDR).

A two-tailed paired t-test showed that the PLD peaked at significantly different latencies between medium (mean ± sd: 188 ms ± 89 ms) and low (224 ms ± 73 ms) discriminability conditions (see Statistics Table 2; Fig. 3A). Since the latency difference between average peaks (140 ms) was close to the difference in average duration of stimuli presentation (SOA) for these two conditions (Δ_SOA_ = 190 ms, Fig. 1B), we investigated whether the PLD difference in latencies was due to the experimental design by calculating the correlation between individual PLD peak latency and average stimuli duration for each participant. No significant correlation was observed (see Statistics Table 3), suggesting that the delay in PLD latencies was not likely due to SOA differences.

If these post-stimulus phase-locking differences found here truly captured the phase reset of the pre-stimulus oscillation, they should show the same frequency dissociation observed in pre-stimulus activity between discriminability conditions. To test this prediction, the PLD at the timepoint of maximum significance of the average traces for the lower-discriminability search conditions (medium: 98 ms, low: 238 ms, stars in Fig. 3A) was plotted across frequencies (Fig. 3B). We observed that the PLD peaked in a significantly higher frequency for the low discriminability task than for the medium one (medium: 9.2 Hz ± 3.0 Hz, low: 14.0 Hz ± 3.6 Hz; see Statistics Table 2), and at similar frequencies as those observed for the pre-stimulus phase opposition. Note that the high discriminability condition is not shown because no timepoint survived the FDR-corrected permutation test in the frequency-band-average analysis (see Fig. 3A).

**Statistics Table 2.**
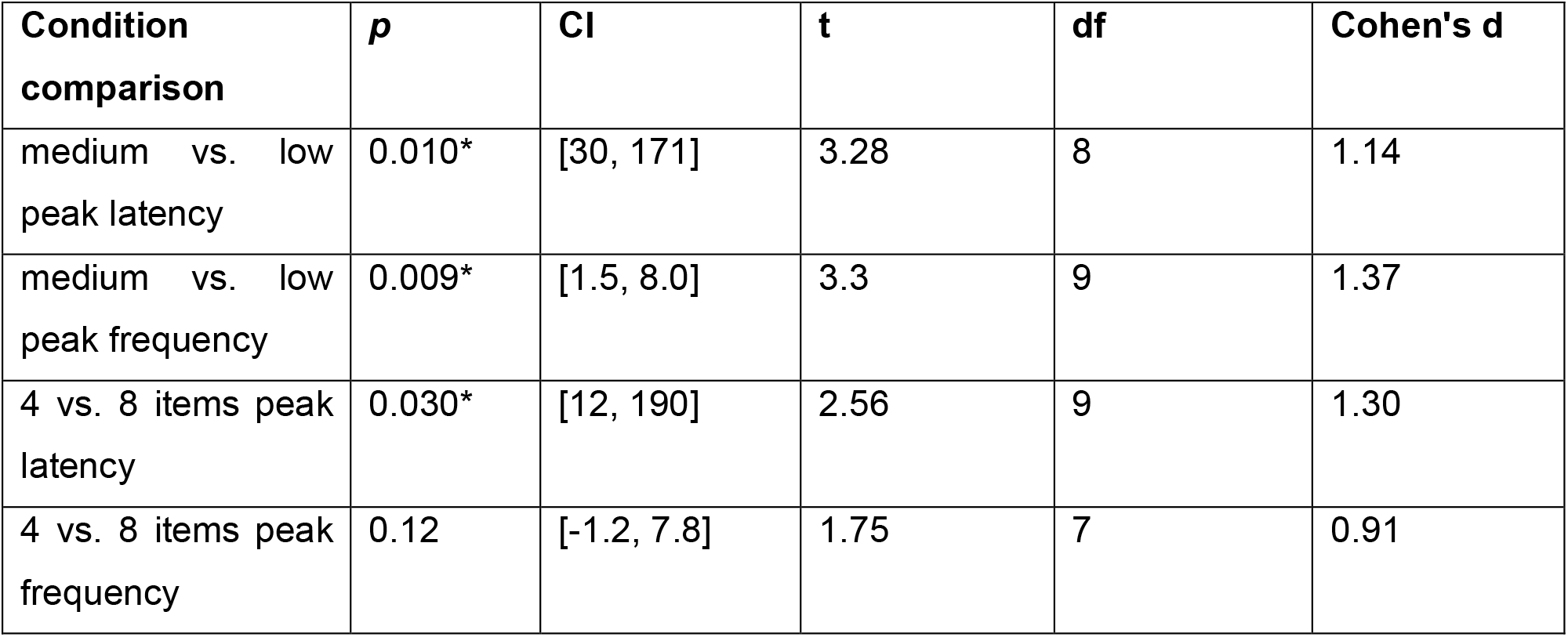
Comparisons between PLD peaks (two-tailed paired t-tests). Same conventions as in Statistics Table 1.

**Statistics Table 3.**
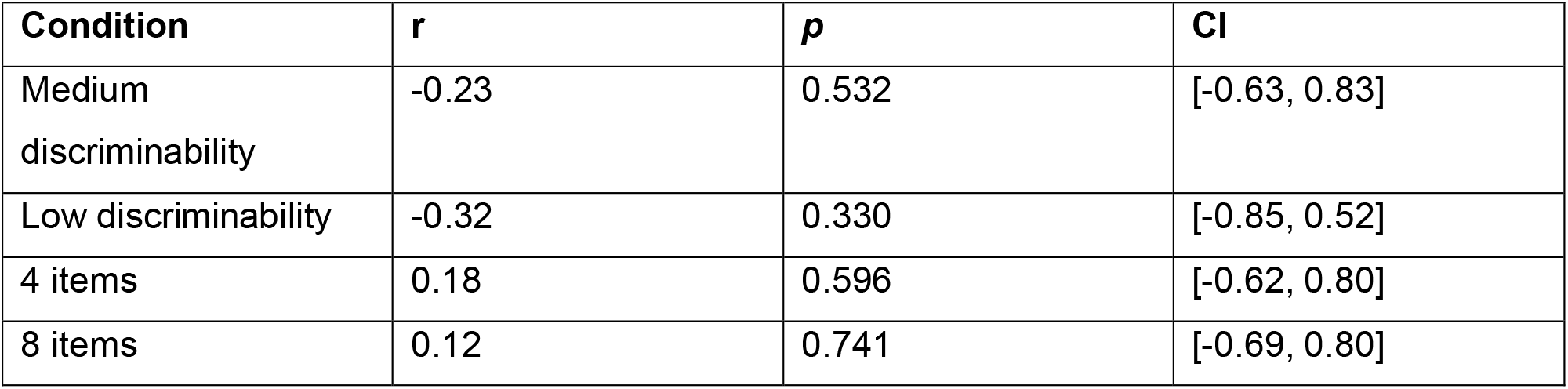
Correlation between individual PLD peak latency and average stimuli duration in each experimental condition. CI: 95% Confidence Interval.

| Condition | *r* | *p* | CI |
| --- | --- | --- | --- |
| Medium discriminability | -0.23 | 0.532 | [-0.63, 0.83] |
| Low discriminability | -0.32 | 0.330 | [-0.85, 0.52] |
| 4 items | 0.18 | 0.596 | [-0.62, 0.80] |
| 8 items | 0.12 | 0.741 | [-0.69, 0.80] |

### Adding items increases the delay of the post-stimulus phase-locking effect

In Experiment 2, the number of items in the search array was manipulated (4 or 8; interleaved). Since participants could not predict the set size of the upcoming trial, we did not test for pre-stimulus frequency effects. Here, we specifically hypothesized that more attentional cycles would be required after stimulus onset if the number of items to process increased, leading to a higher frequency (for the same period of time) and/or a delayed response due to a delay in the post-stimulus phase-locking. We first contrasted the post-stimulus phase-locking between correct and incorrect trials for combined 4- and 8-items conditions. The FDR-corrected permutation test showed that post-stimulus phase-locking value was higher in correct than incorrect trials between 10.2 Hz and 16.9 Hz (FDR p < 1e-8, Fig. 4A). The significant clusters showed a fronto-occipital topographical distribution, similar to the topographies observed in Experiment 1 (Fig. 3A).

**Figure 4.**
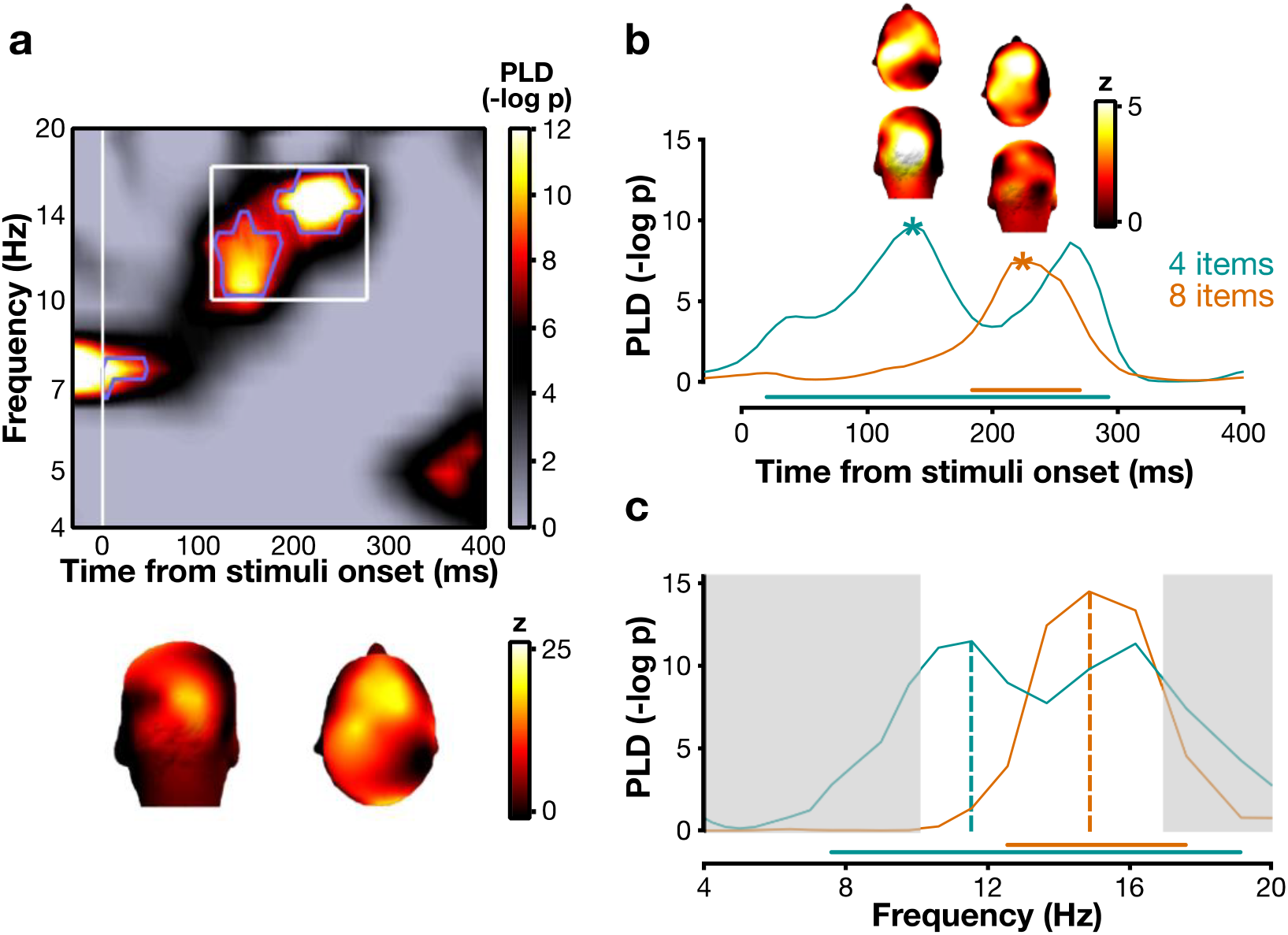
Post-stimulus phase-locking difference between correct and incorrect trials in Experiment 2. (**a**) P value of post-stimulus phase-locking difference (PLD) between correct and incorrect trials for both conditions combined (4 and 8 items), pooled across all electrodes and all participants. Blue contours indicate areas above the FDR threshold (alpha = 10^−7^, corresponding to a p value of 4.4e-9). The bottom topomaps represent the average activity across participants for the time-frequency area within the bounds of FDR-corrected significant blue contours (white rectangle: 114 to 277 ms and 10 to 17 Hz). (**b**) PLD curves plotted after averaging across the frequency range of the significant clusters found in (a). Stars mark the timepoint of maximum significance on each trace (4 items: 137 ms; 8 items: 238 ms). Topomaps represent the PLD at each electrode averaged across participants at the timepoint of maximum significance. (**c**) PLD curves plotted against frequency at the timepoint of maximum significance for the 4-item (teal) and 8-item (orange) condition. On both panels, the horizontal bars along the bottom indicate that the curves surpass the FDR threshold (correction for multiple comparisons). The white area indicates the frequency range of interest, taken from the significant PLD clusters in (a).

We then investigated this effect for each item-number condition separately. We restricted this analysis to the frequency band impacting behavior by averaging the PLD between 10.2 Hz and 16.9 Hz. We found a significant effect of phase-locking on performance in both conditions (4 items: p < 0.01, significant cluster from 20 to 293 ms; 8 items: p < 0.001, significant cluster from 184 to 270 ms; Fig. 4B). Critically, we observed that the 8-items condition was associated with a significantly later peak than the 4-items condition (4 items: 113± 92 ms; 8 items: 223 ± 61 ms; see Statistics Table 2). As in Experiment 1, we calculated the correlation between stimuli duration (SOA) and peak timing to check whether the timing difference could be due to the experimental design. We did not find any significant correlation (see Statistics Table 3), suggesting a dissociation between PLD latencies and SOA. The topographies taken at the points of maximum significance (marked by stars on Fig. 4B) showed that the phase effect is fronto-occipital in both item-number conditions.

Last, we investigated whether the phase-locking difference occurred at different frequencies according to the item condition. We plotted the PLD at the timepoints of maximum significance (4 items: 137 ms; 8 items: 238 ms) against frequency for the two conditions separately (Fig. 4C). Similarly as for Experiment 1 (Fig. 3B), when more items were presented, a trend toward a higher frequency was apparent (4 items: 9.6 Hz ± 4.2 Hz; 8 items: 13.7 Hz ± 2.9 Hz), although this effect was not significant (see Statistics Table 2).

## Discussion

The frequency of attentional sampling reported across the literature shows large discrepancies [4-24]. We propose that part of this heterogeneity is due to inconsistent attentional demands across tasks. The aim of this study was to investigate the link between brain oscillations and attentional demands as manipulated by the discriminability of the target and the set size in a visual search task. We replicated previous findings showing that 7-10 Hz pre-stimulus phase distinguishes correct from incorrect responses [20,32,55-63]. The novelty of the present study lies in the finding that this phase opposition effect is concentrated in rising frequency with degraded discriminability. This link between frequency and discriminability was also found in the post-stimulus phase-locking difference between correct and incorrect search trials. A second independent dataset revealed a similar pattern between the frequency of this post-stimulus phase-locking difference and the number of presented items. Finally, we found significant latency effects of the post-stimulus phase-locking impacting performance both when discriminability was degraded and when distractor number was increased.

We interpret the results under the framework of hierarchical models of visual search. Most theoretical accounts of attentional search propose that lower-level visual areas iteratively communicate with higher-level areas (e.g., [50-53]). Top-down signaling would redirect attention toward candidate items, thus amplifying specific visual neuronal populations consecutively until the target is found. It has been proposed that hierarchical frameworks of visual search involving iterative communication between lower- and higher-level brain regions rest on periodic neural activity in these regions [5,7,20,31,49], possibly involving a causal temporal structuring of visual cortex activity by FEF [70]. If the iterative communication suggested by attentional search models can only process a limited number of items at each iteration, and the time allotted for the search is limited (as in the current experiments), then the frequency should increase with search complexity (lower discriminability or higher set size) to complete more iterations within the same time window. Our results concur with this model, which constitutes the most parsimonious explanation for our findings.

Experiment 1 shows that the less discriminable the target is among the distractors, the higher the frequency of pre-stimulus phase opposition between correct and incorrect searches. This suggests that the neural system can adjust to an optimal frequency, based on goal-directed expectations. Specifically, the frequencies of interest in the high discriminability condition showed no overlap with those in the medium and low discriminability conditions (Fig. 2). However, there was overlap between the medium and low conditions, suggesting that the increase in frequency is not linear across discriminability conditions. This non-linear increase may be related to unequal differences in task demands between conditions, which are difficult to quantify, or to real non-linearity between frequency and task demands. Further investigation is thus needed to distinguish between these two possibilities. A similar frequency modulation was found in the post-stimulus phase-locking effect: the difference in post-stimulus phase coherence between correct and incorrect responses follows increasing frequency with degraded discriminability, as does the pre-stimulus phase effect. This frequency increase is also observed in Experiment 2 (although without reaching the statistical threshold), where trial types were interleaved (i.e., the viewer could not know in advance whether they would search among 4 or 8 items). Together, these results clearly point to a direct link between specific task complexity and brain oscillations. Following hierarchical models’ predictions, this may indicate that the more iterations are performed to find the target, the more neural cycles are generated in the same constrained time window, in line with the hypothesis that more attentional iterations between frontal and occipital regions are implemented through neural oscillations. Our findings are consistent with a previous study, which found that the peak frequency of alpha oscillations involved in a visual discrimination task decreases when the task requires temporal integration relative to temporal segregation (block design) [71]. In the present study, integration could be beneficial for a partially pop-out visual task, such as the high discriminability condition (L vs. +), while temporal segregation would allow processing of more demanding visual searches, potentially requiring sequential processing of stimuli in the search array [25,72,73]. This frequency modulation of pre-stimulus spontaneous oscillations may be controlled by a dynamic top-down signal adjusting to task demands [7,31,34,71,74,75]. One can speculate that a feedback signal gives rise to the lower frequency phase effect found in the high discriminability search condition, thus integrating the search items and facilitating attentional capture by the salient target. A higher frequency in lower-discriminability conditions could, however, enhance segregation between items and contribute to their sequential processing.

The present results show that the phase effects are localized in both low-level (occipital) and frontal regions, which is also coherent with an integration-segregation dissociation mediated by top-down signals. Previous research suggests that the dorsolateral prefrontal cortex (dlPFC) facilitates attentional orienting towards target-like stimuli and comparison between the currently attended stimulus and the target [25,76]. Additionally, oscillations in the frontal eye fields (FEF) are involved in attention shifts [8,25,76]. The theta rhythm was reported to gate and regulate the connections between these anterior and posterior areas [5-7]. Coherent with these previous findings, we observed that the pre-stimulus phase opposition effect is localized over frontal and occipital electrodes for medium and low discriminability, whereas there is minimal frontal contribution in the high discriminability search condition (Fig. 2, bottom). Moreover, the activity in occipital electrodes decreases when there are more items, leaving mainly frontal activity (Fig. 4B). Consistent with a recent study showing minimal involvement of occipital rhythms during an attentional task [77], our observations suggest that frontal areas are more strongly recruited under more complex (difficult) search conditions. Previous literature on exogenous (involuntary) and endogenous (voluntary) attention has shown that exogenous attention relies on bottom-up signals while endogenous attention relies on top-down ones [75,78-84]. Similarly, here, easier search tasks (high discriminability and small set size) may preferentially involve automatic, bottom-up perceptual capture, while more difficult tasks (lower discriminability or large set size) may involve top-down attentional deployment. Indeed, the features that distinguish target (+) and distractors (Ls) in the high discriminability condition are likely to trigger bottom-up attentional capture [85]. Bottom-up signals may be supported by higher frequency bands (hardly recordable using EEG) compared to top-down mechanisms, such as gamma oscillations [86,87]. This limitation can explain the absence of lower-frequency effects in post-stimulus phase-locking in the highly-discriminable, partially exogenous search (Fig. 3A) as well as the absence of frontal involvement in this condition (Fig. 2). Our results therefore suggest that a top-down control tuning the frequency of neural oscillations aids performance in difficult searches.

The results further show that the post-stimulus phase coherence difference between correct and incorrect responses is delayed with increasing complexity (lower target discriminability or larger set size). The mask delay onset, which was adjusted for each participant and condition, could potentially have confounded this effect. Yet, we found no correlation between stimuli duration and peak delay, which argues against this possibility. This is consistent with a previous study using a similar design [20], which found that periodic behavioral modulation was not dependent on stimulus duration during a visual search, and that the onset of the mask did not evoke any performance modulation. We interpret the delayed effect in the framework of two-stage hierarchical models of visual search, i.e., when search tasks are more complex, more iterations between the sensory and the attentional stages are necessary, which is supported by more neural cycles. Such a mechanism implies that the optimal phase reset – leading to the strongest phase-locking – will occur later on average compared to less complex search tasks. Further studies will investigate whether the reset may only affect search while processing the target. Finally, one might notice some discrepancies between effect localizations (Fig. 3): both the low and medium discriminability conditions showed an occipital effect, but while the low condition showed also a frontal activation, the medium condition showed a parietal one. This difference might be explained by the involvement of different brain areas, hence explaining the different delays of the effect between the two conditions. It might also come from differences in dipoles within similar brain areas, which can project differently onto the sensors. Further investigation is thus warranted.

This study contributes to filling the missing experimental evidence for the current models of attentional search and provides novel support for a link between brain oscillations and attentional demands, which may be a factor to explain the frequency variability observed in the literature. Our results suggest that brain oscillations support attentional sampling, which may correspond to the iterations posited by biologically-plausible computational and cognitive models of attentional search. Importantly, the pre-stimulus modulation of frequency, accompanied by post-stimulus frequency and latency modulations in the same direction, indicate a capacity for low-frequency oscillations (< 20 Hz) to structure the temporal dynamics of neural activity and aid top-down, attentional control for maximally efficient visual processing.

## Materials and Methods

### Participants

Thirteen participants (6 women) were recruited for Experiment 1. One was excluded from analyses because they were not able to maintain fixation. Thirteen different participants (5 women) were recruited for Experiment 2. Two were excluded due to poor EEG signal quality. All participants (age between 18 and 35 years) had normal or corrected-to-normal vision and declared no previous history of neurological disorders or diagnoses. All participants provided written informed consent and received monetary compensation for their time. The study followed the principles prescribed by the declaration of Helsinki and was approved by the ethics committee “CPP Sud-Ouest et Outre-Mer I,” under protocol number 2009-A01087-50.

### Apparatus and Stimuli

Both experiments were conducted in a dark room, with the participant sitting 57 cm away from a CRT monitor (800×600 pixels; refresh rate: 140 Hz), their head positioned using a chinrest. The stimuli were red letter-like symbols (oriented randomly 0°, 90°, 180° or 270° from vertical) and masks (squares barred by a cross of the same size as the stimuli) presented away from a central fixation dot on a uniform gray background, created under the MATLAB software (The MathWorks, www.mathworks.com, version R2016b) using the Psychophysics Toolbox [88-90].

In Experiment 1, the shape of the stimuli was manipulated. Four stimuli (1.8° x 1.8° visual angle) were presented simultaneously, either on the lower right or lower left quadrant of the screen (randomly), at a fixed eccentricity (8°). In the high discriminability condition, the distractors were L shapes and the target was a + shape. In the medium condition, the distractors were L shapes and the target was a T shape. In the low discriminability condition, the distractors were distorted L shapes with the horizontal bar slightly shifted up so they resembled a T, and the target was a T shape (Fig. 1A).

In Experiment 2, the set size was manipulated. The distractors were always L shapes and the target a T shape. In the 4-items condition, four stimuli were presented at 8° of eccentricity (one at each of the cardinal positions around fixation), and in the 8-items condition, eight stimuli were presented: one at each cardinal position and one at each diagonal position around fixation (Fig. 1A).

### Experimental Procedure

#### Behavioral pre-test for Experiment 1

To assess the discriminability of items in the attentional search task for each condition, the participants from Experiment 1 (not Experiment 2) first performed a behavioral pre-test at the beginning of the experimental session testing the three different stimuli shapes (L vs. +, L vs. T or distorted-L vs. T) and the two set sizes (4 or 8 items). Trials in this behavioral pre-test were blocked by shape condition (104 trials per shape), with the two set sizes pseudorandomly interleaved in every block (52 trials per set size in each block). Participants were instructed to report as fast as possible whether the target was present or absent in the array, and the stimuli stayed present on the screen until the participant pressed a response key (left arrow for target present, and right arrow for absent). A t-test on median reaction times shows a weak but significant difference between 4 and 8 for L vs. + items (4 items: 548 ± 46 ms, 8 items: 580 ± 48 ms, median ± 95% CI, p = 0.0058, CI = [-29,-6], t(10) = -3.50). There is a greater difference between 4 and 8 items with L vs. T (4 items: 591 ± 52 ms, 8 items: 759 ± 75 ms, median ± 95% CI, p < 0.0001, CI = [-144,-71], t(10) = -6.51) and with distorted-L vs. T (4 items: 1696 ± 224 ms, 8 items: 2426 ± 408 ms, median ± 95% CI, p < 0.0001, CI = [-1208,-672], t(10) = -7.81). Crucially, the reaction times are significantly more increased as discriminability decreases (L vs. +: Δ_RT_ median ± 95% CI = 22 ± 10 ms; L vs. T: 131 ± 32 ms; distorted-L vs. T: 1052 ± 236 ms; t-test of L vs. + against L vs. T: p < 0.001, CI = [-129,-51], t(10) = -5.18; L vs. + against distorted-L vs. T: p < 0.0001, CI = [-1192,-653], t(10) = -7.63; L vs. T against distorted-L vs. T: p < 0.0001, CI = [-1089,-576], t(10) = -7.23). We therefore label the L vs. + condition as a high discriminability condition, the L vs. T as medium and the distorted-L vs. T as low discriminability. We selected the stimuli shape from the medium discriminability condition (L vs. T) for Experiment 2, in which we then manipulated set size.

#### EEG sessions

In both experiments, at each trial, participants were asked to fixate a central dot. Their eye position was verified a posteriori using the electro-oculography (EOG) signal, and only trials with no blink nor saccade were kept for the analyses (mean ± sd: 158 ± 50 rejected epochs). Participants began each trial by pressing a key. After a random temporal jitter of 1.5 to 2.5 seconds, the stimuli appeared on the screen for the duration of the Stimulus-mask Onset Asynchrony (SOA) defined for each individual, and were directly followed by masks, for a total duration of 500 ms. The SOA was titrated for each participant and each condition in a preliminary staircase block with an adaptive staircase procedure [91] to obtain an average performance level of 70% correct responses. The SOA was re-adjusted after each block if necessary, to maintain this level throughout the experiment. The average SOA across all participants was 40 ± 5 ms (SD) for the high discriminability, 82 ± 13 ms for the medium, and 271 ± 26 ms for the low discriminability condition in Experiment 1; and 69 ± 43 ms for the 4-items condition and 129 ms ± 78 for the 8-items condition in Experiment 2. The target was present randomly on half of the trials. Participants were asked to report whether the target was present or absent as accurately as possible, without a time limit. Participants reported the presence or absence of the target using a keyboard (right arrow key for present and left arrow key for absent). All participants used their right hand.

In Experiment 1, participants performed 6 blocks (2 of each discriminability condition) in pseudorandomized order across participants, with 250 trials per block, leading to 500 trials per condition in total. In Experiment 2, the set size conditions (4 items or 8 items) were pseudorandomly interleaved within the blocks. Participants performed 30 blocks with 40 trials per block, leading to 600 trials per condition in total. The total experiment duration varied between 2 and 2.5 hours to account for the EEG setup, the behavioral pre-test (for Experiment 1), the staircase procedure (for both Experiments 1 and 2), and breaks.

### Behavioral analyses and statistics

The behavioral dependent variable was participants’ percentage of correct responses, which was aimed to be maintained at 70% to ensure that participants were not guessing the answer and that there was a balanced ratio of correct and incorrect trials. A trial was labeled as correct if the participant correctly identified that a target was present or correctly reported that the target was absent. Trials which were rejected in EEG pre-processing were also deleted from the behavioral analysis. Conditions were compared using a repeated-measures one-way ANOVA in Experiment 1 (three conditions) and two-tailed t-tests (post-hoc in Experiment 1). The present study was not designed to measure differences between target-present and target-absent conditions, and therefore does not include sufficient numbers of trials in these constrained cases [68]. Future experiments are needed to investigate this difference.

In the pre-test for Experiment 1, the principal dependent variable was the reaction time (measured between stimulus onset and the participant’s response) as a function of the number of stimuli in the search array (4 or 8). This was measured for each level of discriminability (high, medium and low). Only the trials in which participants gave a correct response were included in the analysis. This pre-test enabled us to compute the difference between median reaction times (Δ_RT_ or slope) for 4 vs. 8 items, and thus to measure how strongly the number of distractors affected performance, independently for each shape condition. The reaction times when 4 or 8 items were presented, and the slopes for each shape condition, were compared using pairwise (2-by-2) t-tests.

### EEG acquisition and pre-processing

EEG was recorded using a 64-electrode ActiveTwo Biosemi system. Two extra electrodes, Common Mode Sense (CMS) and Driven Right Leg (DRL) were used as reference and ground. They were placed on the participant’s face, with CMS on the left cheek and DRL on the right. Eye movements were recorded with EOG using three additional electrodes: one on each side of the eyes (to record horizontal eye movements) and one under the right eye (to record vertical movements).

All EEG analyses were performed under MATLAB using the eeglab toolbox [92] and custom scripts. Numerical computations were partly performed on the S-CAPAD platform, IPGP, France. The data were downsampled to 512 Hz, re-referenced to the average over all electrodes, and epoched from -1500 to +1000 ms relative to search array onset. The data were cleaned by visual inspection to remove artifacts: large spurs of high-amplitude, high-frequency activity in one or more electrodes (muscle contractions), blinks and saccades, as made evident by a sharp depolarization and repolarization in the EOG (eye movements), slow drifts of large amplitude in more than 3 electrodes. In addition, electrodes were interpolated using spherical spline interpolation if they read noisy signal for more than 7 % of the trials. On average, 134 ± 20 (mean ± sd) epochs were rejected per participant (∼9 %) in Experiment 1 and 182 ± 80 epochs in Experiment 2 (∼15%). Epochs were then labeled according to performance (correct or incorrect) for each condition.

### Phase and amplitude analyses

Time-frequency decomposition was performed using Morlet wavelets on single trials [93]. The wavelets’ length evolved logarithmically from 3 to 6 cycles between 4 and 20 Hz. The signal was expressed at every time-frequency point as a complex vector. Inter-Trial phase Coherence (ITC; otherwise called phase-locking) was computed by averaging the complex exponentials over all trials of a given condition for each time-frequency point, electrode and participant. This analysis was done separately for correct and incorrect trials in each condition. To account for the imbalanced ratio of correct/incorrect trials (70%/30%), the correct trials were randomly subsampled to attain the same number as the incorrect trials (repeated 100 times).

To analyze whether the pre-stimulus phase between correct and incorrect trials differed, Phase Opposition Sum (POS) was computed at each time-frequency point:

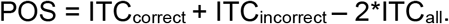

A low POS indicates that the two conditions (correct and incorrect) have random phase distributions, whereas a high POS means that phase-locking is strong in both conditions. The measure of POS accounts for uniform distributions of phases: to ensure that the phase was truly opposed between correct and incorrect trials, we checked for circular uniformity (Hodges-Ajne test) of the phase angles (see [68,69]) across all trials (characteristic of a spontaneous signal) at the point of maximum significance at sensor Oz, because it is activated in the 3 conditions (high, medium and low discriminability; see Fig. 2), and ensured that no instance of non-uniformity was found (high discriminability: *p* = 0.242, kappa = 0.3; medium: *p* = 0.854, kappa = 0.148; low: *p* = 0.92, kappa = 0.037). Therefore, a strong phase-locking in one condition presupposes a phase-locking in the other condition in opposite phase, and thus a high POS [52].

We also measured post-stimulus Phase-Locking Difference (PLD) between correct and incorrect trials, defined as: PLD = ITC_correct_ - ITC_incorrect_. A high PLD reveals a strong phase-locking for correct trials and a weak phase-locking for incorrect trials, meaning that one precise phase for the reset aided performance (correct trials), and trials resetting away from this ideal phase led to low performance on average (incorrect trials). As mentioned in the introduction, the term “reset” is used here purely to refer to the mathematical description of the resulting signal.

Using the time-frequency decomposition by Morlet wavelet described above, the amplitude for each time-frequency point was also computed by taking the length of each complex vector. Baseline correction was applied by subtracting the mean value from -0.4 to -0.1 s from stimulus onset. The same subsampling procedure as for the ITC calculation was performed.

The POS and PLD significance was tested using a permutation procedure. Following the null hypothesis according to which there is no difference between correct and incorrect trials, we compared the averaged phase-locking values to shuffled surrogates of the data within each participant. The surrogates were generated by shuffling “correct” and “incorrect” trial labels while keeping the original number of incorrect trials and subsampling correct trials as described above. This procedure was repeated 10,000 times to obtain a null distribution. The POS and PLD were converted to a z-score, by subtracting the mean of the null distribution (shuffled surrogate) from the real POS or PLD, and dividing by the null distribution’s standard deviation. This yielded a z-score for each participant, which were then combined using the Stouffer method [68,94] to obtain a group-level z-score. This z-score was expressed as a p-value using the cumulative distribution function. Multiple comparisons were corrected using the False Discovery Rate (FDR) procedure, in which the p-values associated with each time-frequency point are ordered and thresholded [95], with a threshold α value of 10^−7^. To test for the presence of outliers in the data, a leave-one-out procedure was performed wherein one participant at a time was removed from the pool.

These analyses were performed from -0.4 to 0 s (around search array onset) for the POS and 0 to 0.4 s for the PLD. The same statistical procedure was applied to amplitude. Finally, individual participants peak values POS, and PLD were compared across experimental conditions. Participants that did not exhibit any significant POS or PLD peak individually were excluded from the comparison analyses (in Experiment 1, three participants were excluded from the latency comparison and two from the frequency comparison, resulting in a total n of 9 and 10, respectively; in Experiment 2, one participant was excluded from the latency comparison and three from the frequency comparison, resulting in a total n of 10 and 8, respectively).

## Data Availability

The datasets generated during and/or analyzed during the current study are available from the corresponding author upon reasonable request.

## Acknowledgements

This project has received funding from the Agence Nationale de la Recherche (ANR) - Deutsche Forschungsgemeinschaft (DFG) programme (grant agreement No J18P08ANR00 – Laura Dugué), the ANR programme (grant agreement No ANR-19-NEUC-0004 – Rufin VanRullen), and the Université de Paris IDEX doctoral programme (grant No J18I08IDEX13 – Garance Merholz).

## Author contributions

Conceived and designed the experiments: LD, RV. Performed the experiments: LD. Analyzed the data: GM, LG, LD. Prepared the figures: GM. Reviewed and edited the figures: GM, LG, LD. Wrote the first draft of the manuscript: GM. Reviewed and edited the manuscript: GM, LG, LD, RV.

