## Supplementary Material for "Periodic attention operates faster during more complex visual search"

### Supplementary material for:
### Periodic attention operates faster during more complex visual search

Garance Merholz[1]*, Laetitia Grabot[1], Rufin VanRullen[2], Laura Dugué[1,3]

[1]Université de Paris, INCC UMR 8002, CNRS, F-75006 Paris, France

[2] Centre National de la Recherche Scientifique, CerCo Unité Mixte de Recherche 5549, Université de Toulouse, Toulouse F-31052, France

[3]Institut Universitaire de France (IUF), Paris, France

*

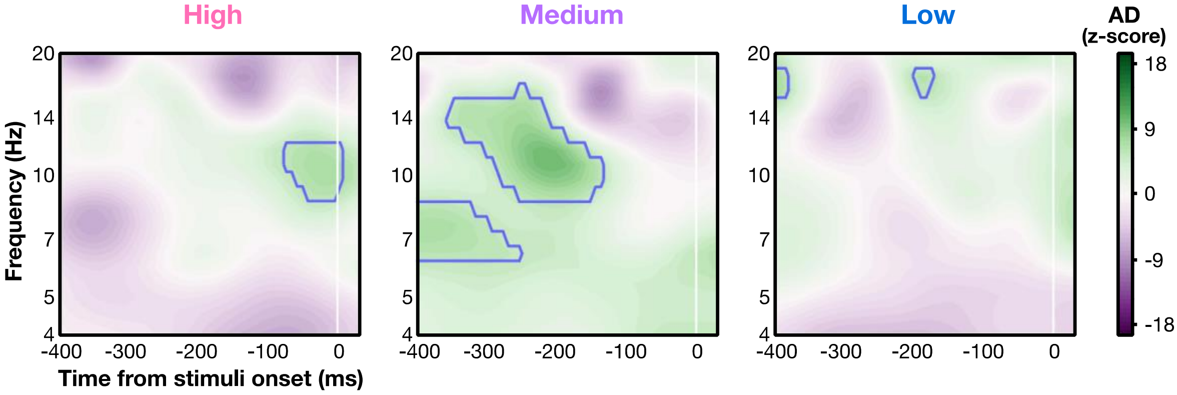


***Figure S1. Pre-stimulus amplitude difference between correct and incorrect trials in Experiment 1.*** *Z-score of pre-stimulus amplitude difference (AD) between correct and incorrect trials for the high, medium and low discriminability conditions, combined across all electrodes and all participants. Blue contours indicate areas above the FDR threshold (alpha = $10^{-7}$, corresponding to p values of 2.2e-9, 1.8e-8 and 8.2e-10, respectively).*
